# Purinergic signalling and calcium dynamics: potential drivers in the onset of coordinated intestinal motility in human fetal development

**DOI:** 10.1101/2025.04.17.649306

**Authors:** Benjamin Jevans, Atchariya Chanpong, Luca Peruzza, Silvia Perin, Ben Cairns, Parth Tagdiwala, Pieter Vanden Berghe, Grant W Hennig, Conor J McCann

## Abstract

**Background and aims:** Intestinal motility relies on inputs from multiple cells within a complex neuromuscular syncytium located in the gut wall. While it is known that motility is dependent on the development of motor patterns which govern coordinated contractile activity within the gut wall, knowledge regarding the onset of coordinated motor activity in humans is still lacking. This study assessed the emergence of coordinated motor patterns, and the molecular mechanisms underpinning developing motility, in the human fetal gut.

**Methods:** Human fetal gut samples (obtained via the MRC-Wellcome Trust Human Developmental Biology Resource-UK) were characterised by live imaging, spatiotemporal mapping, immunohistochemistry and RNAseq.

**Results:** Human small intestinal samples displayed the presence of key cell types including enteric neurons, interstitial cells of Cajal, platelet-derived growth factor receptor alpha positive (PDGFRα+) cells and smooth muscle at post conception week (PCW) 12. Between PCW12 and PCW16, functional assessment revealed a marked increase in the velocity (p= 0.0341) of propagating contractions. Subsequently, between PCW16 and PCW20 the number of contraction initiation sites reduced drastically (p=0.0053), enabling the emergence of long-distance propagating contractions. Expression analyses showed the development of coordinated motor activity was coincident with increased expression of various genes involved in calcium and purinergic signalling pathways.

**Conclusions:** These findings provide the first direct mechanistic evidence of the temporal development of coordinated contractile activity in the human fetal intestine, highlighting the role of calcium dynamics, purinergic signalling and interstitial cells in early stages of human motility development, potentially informing an improved understanding of the pathogenesis of gut motility disorders.

## Introduction

Gastrointestinal (GI) disorders have a worldwide prevalence of approximately 40%^1^, placing a huge burden upon healthcare systems and significantly affecting patient quality of life. For the vast majority of these conditions there remains no cure and a paucity of treatment options^2, 3^, often due to a limited understanding of the underlying physiological processes.

Correct functioning of the GI tract is crucial, with numerous cell types working in close coordination to fulfil a number of functions, chiefly the digestion of ingested food into absorbable nutrients and egestion of waste. Movement along the intestine is achieved by peristalsis: a series of coordinated muscle contractions and relaxations. This coordination relies on the highly orchestrated development of the enteric neuromusculature, with inputs from both the enteric nervous system (ENS, the intrinsic innervation of the gut) and specialised interstitial cells, including interstitial cells of Cajal (ICC) and platelet-derived growth factor receptor alpha positive (PDGFRα+) cells^4, 5^. These are electrically coupled to smooth muscle cells (SMCs) in the SIP (**<u>S</u>**MCs, **I**CC, **<u>P</u>**DGFRα+ cells) syncytium^6^. Constant communication between the SIP syncytium and ENS allows the GI tract to conduct its functions with almost exclusive autonomy from the central nervous system. The vital contributions of each system are evidenced by the severe impacts on GI motility observed in mouse models exhibiting mutations in genes for either ENS development^7^ or ICC^8^ and PDGFRα+ cell function^9^. Interestingly, despite their different cellular origins, diseases affecting the ENS also impact density and distribution of both ICCs and PDGFRα+ cells, demonstrating the synergistic nature of the SIP and ENS^10^.

Recent spatiotemporal analysis of gene expression in the human fetal small intestine revealed the presence of neuronal progenitors as early as post-conception week (PCW) 8 and ICC and PDGFRα+ cells at <PCW12^11^ suggesting the early presence of each cell type within the developing human intestine. Interestingly, *in utero* studies have detected gastric contractions as early as PCW14^12^ and *ex vivo* analysis in isolated human fetal jejenum identified myogenic ripples as early as PCW13^13^. Further, recent investigations have demonstrated that the onset of compound electrical activity in the human colon occurs between PCW14-16^14^ and that neurally-mediated contractile activity can be identified in the jejunum at PCW18^13^. However, our understanding of how coordinated GI motility develops, in line with the growth of the intestine and the key molecular pathways involved in establishing early motility patterns in humans, remains sparse.

Here, we show that development of coordinated motor patterns in the human fetal small intestine occurs between PCW12-20. We demonstrate that development of this coordinated behaviour coincides with significant alterations in gene expression patterns relating to calcium dynamics and purinergic signalling, suggesting a role of interstitial cells in the development of coordinated activity in the early fetal gut. Hence, we propose that the early second trimester period is crucial for the spatiotemporal organization of coordinated neuromuscular activity in the human intestine.

## Methods

### Human sample collection

Human small intestine samples were collected, under informed ethical consent, through the Joint MRC/Wellcome Trust Human Developmental Biology Resource with Research Tissue Bank ethical approval (08/H0712/34+5 and 08/H0906/21+5). A total of 18 individual samples were used (10 female and 8 male samples, **Supplementary methods table 1**).

### Imaging Small Intestine Contractility

Segments of small intestine were harvested from between 2-8 cm above the caecum and pinned in a Sylgard-lined dish containing oxygenated Krebs solution (composition: NaCl 120.9 mM, KCl 5.9 mM, MgCl_2_ 1.2 mM, Glucose 11.5 mM, NaHCO_3_ 14.4 mM, NaH_2_PO_4_ 1.2, CaCl_2_ 2.5 mM) at 37°C. The mesentery was excised by sharp dissection and the segments left to equilibrate for 30 mins prior to imaging. Images were captured using µManager software (FIJI) via a USB camera (DMK 42AUC03, The Imaging Source, Germany) connected on a custom designed stereo mount with 10:1 macro zoom manual lens (MLH-10X, The Imaging Source, Germany). The contractile activity of the intestinal segments was recorded at 2 frames per second for 5-10 minutes. Images were processed and spatiotemporal maps (ST Map) created, and analysed, using custom-written software (Volumetry G9j: GWH). Please see supplementary M&Ms for more information on specifics on ST Map construction and analysis.

### RNAseq analysis

RNA was extracted from human fetal small intestinal samples collected at PCW12, PCW16 and PCW20. For each age, n=3 samples were used. Briefly, samples were homogenised in appropriate volumes of Trizol (Qiagen, U.K.) and RNA isolated following addition of chloroform (VWR, Pennsylvania, U.S.A.). The aqueous phase was processed using an RNEasy Mini Kit (Qiagen, U.K.) following the manufacturer’s instructions. DNase I (Qiagen, The Netherlands) was used to remove DNA contamination. Sequencing libraries were generated by UCL Genomics (London, U.K.) using the KAPA RNA HyperPrep Kit with RiboErase (HMR, KK8561, Roche, Basel, Switzerland) according to the manufacturer’s instructions and sequenced at 800pM on a NextSeq 2000 (Illumina, San Diego, US), using a 57bp paired-read run with corresponding 8bp unique dual sample indexes and 8bp unique molecular index. RNA concentration and integrity were quantified using a Nanodrop and Agilent’s 4200 Tapestation (Standard Total RNA assay): the average RIN number for PCW12, PCW16 and PCW20 was 7.0, 5.0 and 6.3, respectively and average RNA concentration for each age was 74.5, 95.7 and 84.9ng/µL, respectively. See Supplementary materials and methods (M&Ms) for details on analysis parameters.

### Immunofluorescence investigations

Human small intestinal samples were cleared of mesentery, fixed in 4% paraformaldehyde (PFA, Sigma Aldrich, U.K.), snap-frozen in gelatin and sectioned (25µm) using a Leica CM1900 UV Cryostat (Leica Microsystems, U.K.). Following thawing at room temperature (RT), slide-mounted cryosections were post-fixed in 4% PFA. Slides were washed and blocked in 1% bovine serum albumin (Sigma Aldrich, U.K.) and 0.1% Triton X-100 (Sigma Aldrich, U.K.) in 1XPBS at RT. Primary antibodies (**Supplementary methods table 2**) were diluted in blocking solution and applied overnight at 4°C. Secondary antibodies (**Supplementary methods table 2**) were diluted in blocking solution and applied for 2 hours at RT. Vectashield (Hard Set, Dako, U.K.) was used to cover-slip slides, which were stored at 4°C until imaging. See Supplementary M&Ms for quantification methods.

### Statistical analysis

Analysis of motility data and immunofluorescent stains was conducted on Graphpad. Bar graphs are plotted as mean data values +/-SEM. Statistically significant changes between the three ages were assessed using ANOVA, with p values <0.05 taken as significant.

## Results

### Human small intestinal tissue contains key functional cell types at PCW12

GI motility is controlled by a complex interplay involving the ENS and SIP syncytium. Earlier reports have demonstrated the presence of neurons in the human colon from PCW12^14^. We verified similar expression in human small intestinal samples using the pan-neuronal marker TUBB3 (**Fig. 1** and **Supp. Fig. 1A-C**). TUBB3+ cells were apparent within both the myenteric (**Fig. 1A**, arrows) and submucosal plexuses (**Fig. 1A**, arrowheads) at PCW12. These structures appeared largely unchanged at PCW16 (**Fig. 1B** and **Supp. Fig. 1B**) and PCW20 (**Fig.1C** and **Supp. Fig. 1C**). However, at PCW20, the ganglia within the myenteric plexus appeared more discrete and dense (**Fig. 1C**, arrows and **Supp. Fig. 1A-C**).

**Figure 1:**
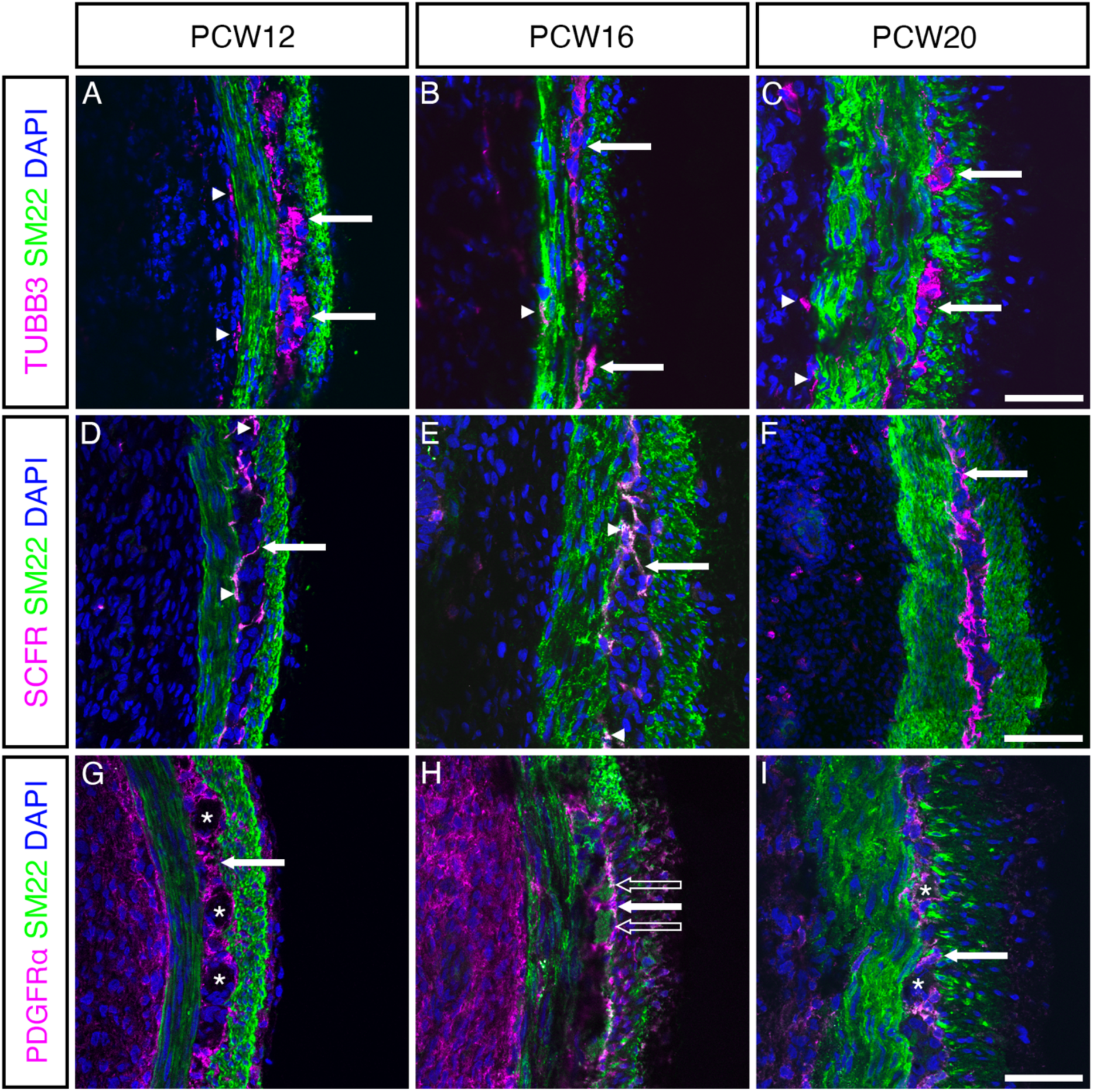
Neurons, ICC and PDGFRα+ cells are present in the small intestinal wall from PCW12. (A-C) Representative merged fluorescent images showing SM22 (green), DAPI (blue) and TUBB3 (magenta) expression in cryosections of human small intestine at PCW12, PCW16 and PCW20. TUBB3+ neurons were clearly identified at the level of the myenteric (arrows) and submucosal (arrowheads) plexuses. (D-F) Representative merged fluorescent images showing SM22 (green), DAPI (blue) and SCFR (magenta) expression across all time points. SCFR+ ICCs were identified at each timepoint at the level of the myenteric plexus with the presence of filamentous SCFR+ projections (arrows). At PCW12 and PCW16 SCFR+ cells that co-express SM22 could also be identified (arrowheads). (G-I) Representative merged fluorescent confocal images showing SM22 (green), DAPI (blue) and PDGFRα (magenta). PDGFRα+ expression was observed across all time points. The majority of PDGFRα cells were found between the circular and longitudinal muscle layers (arrows). PDGFRα+ cells often appear to envelope other structures, including several which did not stain positive for SM22 (G, I, asterisks) as well as some SM22+ structures (H, hollow arrows). Scale bars 50µM. **Refer to Supp. Figs. 1-3 for individual split channel images from which the merged panels shown in Fig. 1 were generated.**

We also detected stem cell factor receptor positive (SCFR+) ICC in the human small intestine (**Fig. 1D-F** and **Supp. Fig. 2**). At PCW12, PCW16 and PCW20, SCFR+ cells were predominantly located at the interface between the circular (CM)/longitudinal muscle (LM) layers at the level of the presumptive myenteric plexus, with occasional projections traversing this space (**Fig. 1D-F**, arrows). At PCW12 and PCW16, SCFR+ cells were observed co-labelled with the smooth muscle marker SM22, although this was less apparent at PCW20 (**Supp. Fig. 2**) whereby distinct SCFR+ staining of the ICC-MY network appeared more robust. PDGFRα+ cells were also found within human small intestinal tissue at PCW12, PCW16 and PCW20 (**Fig. 1G-I** and **Supp. Fig. 3**). Similarly to SCFR, cells displaying robust PDGFRα+ expression were frequently detected at the level of the myenteric plexus (**Fig. 1G-H**, arrows and **Supp. Fig. 3**). At PCW12, PDGFRα+ staining was distinct from SM22+ staining, but at PCW16 and PCW20 PDGFRα+ and SM22+ staining was often colocalised (**Supp. Fig. 3**) with PDGFRα+ cells often encircling other, unlabelled, presumptive ganglia, structures (**Fig. 1G, I**, asterisks). However, occasionally, these structures were found to be SM22+ (**Fig. 1H**, unfilled arrows). Finally, we observed diffuse PDGFRα+ signals within the submucosa (**Supp. Fig. 3G-I**, ‘SM’ unfilled arrowheads) and at the serosal boundary (**Supp. Fig. 3G-I**, ‘S’ hollow arrowheads) at each developmental timepoint investigated.

### Contractility in the human small intestine becomes increasingly coordinated between PCW12 and PCW20

Having confirmed that cells of the ENS and SIP syncytium were present at PCW12, with morphological evidence of maturation between PCW12 and PCW20, we next sought to characterize SI motility at these timepoints (see Methods and **Supp. Fig. 4**). Spatio-temporal (ST) mapping and quantification of diameter, frequency and velocity parameters (**Fig. 2A-C**) revealed distinct motility patterns in both PCW16 and PCW20 intestinal segments compared to PCW12 samples. The average outer diameter of intestinal preparations doubled in size between PCW12 and PCW20 (PCW12: 1.05 ± 0.06mm; PCW16: 1.71 ± 0.1mm; PCW20: 2.26 ± 0.25mm, p= 0.0017, **Fig. 2A-D** and **Supp. Fig. 5A, B**).

**Figure 2:**
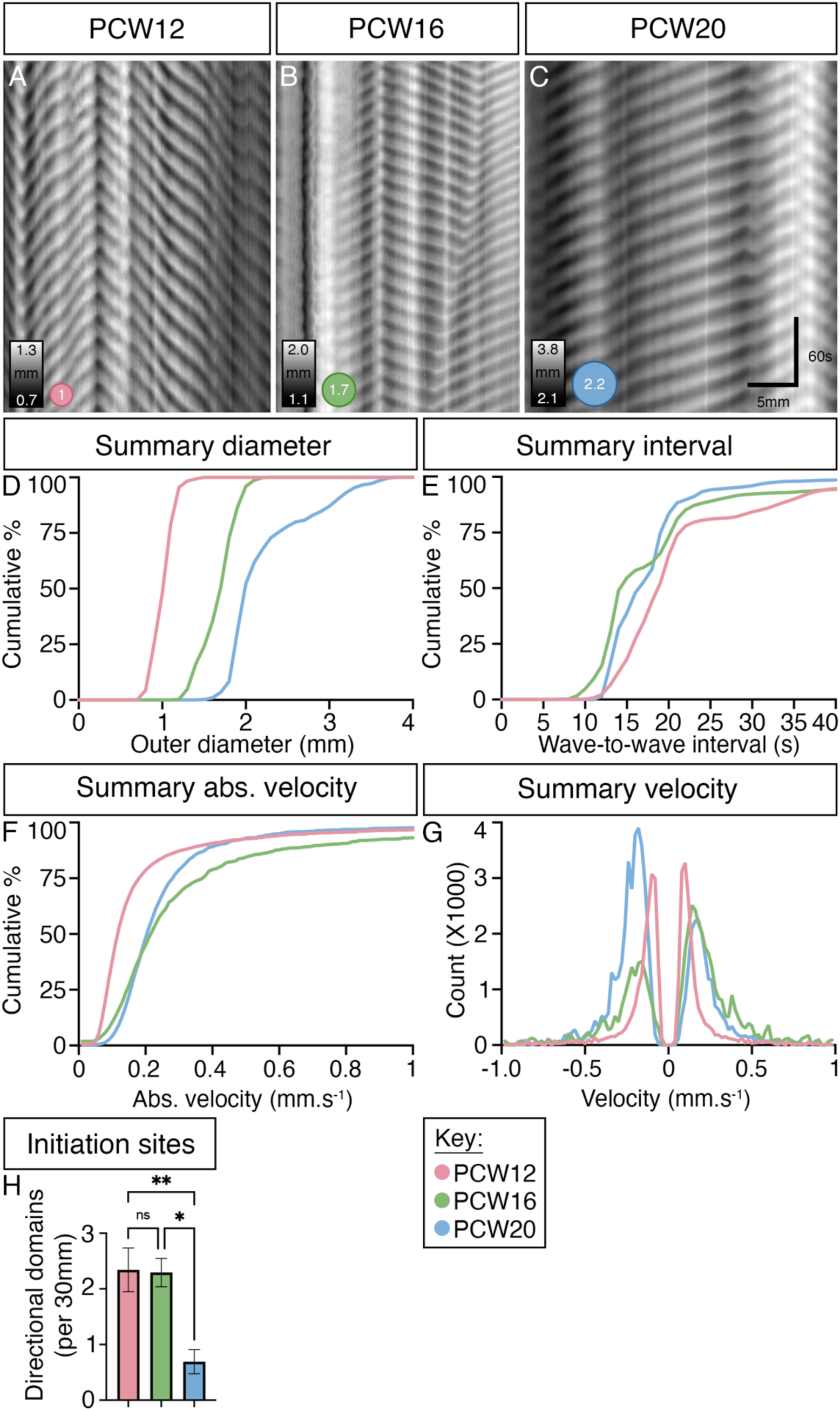
Changes in contractile characteristics of small intestinal motility with age. (A-C) Representative ST map examples of small intestinal motility at ages PCW12 (A: pink), PCW16 (B: green) and PCW20 (C: blue). The median diameter at each PCW (n=4 for each group) is displayed diagrammatically as a circle in the lower left-hand corner of each ST Map. Note the different diameter ranges for each ST map (rectangular scalebar of diameters: grayscale). (D) Cumulative frequency histograms of intestinal diameter at each of the three PCWs showing a significant increase in overall outer diameter between PCW12 to PCW16, with a slower increase between PCW16 to PCW20 (**refer to Supp. Fig. 5A for statistical comparisons**). (E) The overall frequency of contractions was fastest (shortest interval) at PCW16 and slowest (longest interval) at PCW12 (**refer to Supp. Fig. 5B for statistical comparisons**). (F) The absolute velocities of propagating contractile waves were significantly slower (∼50%) in intestinal segments from PCW12 compared to PCW16 (**refer to Supp. Fig. 5C for statistical comparisons**). There was no further difference between segments from PCW16 and PCW20. This suggests, like for other parameters, that intestinal segments from PCW12 are not fully developed to allow transmission of excitation along the gut wall at the same level compared to PCW16 and PCW20. (G) The overall proportion and velocity of orally and anally-propagating contractions was mixed, depending on where and how many initiation sites were present along each intestinal segment. (H) Summary data showing the number of initiation sites per age group.

Repetitive propagating contractions, seen in ST Maps as a series of sloped black and white stripes (**Fig. 2A-C**), appeared to have a regular frequency and velocity. However, analysis showed that intestinal segments from PCW12 tended to have a slightly slower average frequency (longer interval between successive waves, PCW12: 22.58 ± 6.43s; PCW16: 19.8 ± 7.54s; PCW20: 18.3 ± 3.67s, p= 0.8852, **Fig. 2E**, and **Supp. Fig. 5C, D**) and significantly slower median propagation velocity compared to PCW16 and PCW20 (PCW12: 0.12 ± 0.02mm.s^-1^; PCW16: 0.2 ± 0.03mm.s^-1^; PCW20: 0.21 ± 0.01mm.s^-1^, p= 0.0341, **Fig. 2F, G** and **Supp. Fig. 5E, F**).

Surprisingly, analysis of the number of unidirectional domains/initiation sites showed a dramatic and significant decrease at PCW20 (PCW12: 2.34 ± 0.39, PCW16: 2.29 ± 0.25, PCW20 0.69 ± 0.22, p=0.0053, **Fig. 2H**). Indeed, in 3 of 4 PCW20 preparations, all contractions propagated in only one direction across the preparations, demonstrating that slow waves generated at initiation sites could be entrained and propagate over much larger distances compared to earlier weeks.

Taken together, these data demonstrate that there are 2 defined phases to the maturation intestinal motor behaviour: the first occurs between PCW12 and PCW16 where the velocity and frequency of propagating contractions increases markedly, and the second between PCW16 and PCW20 where the number of contraction initiation sites is drastically reduced allowing contractions to propagate long distances without interference.

### Increasing contractile coordination with age between PCW12 and PCW20 coincides with increasing ICC network maturity

Given the marked delay in the onset of coordinated activity, despite immunohistochemical detection of all enteric and SIP cell types in the correct anatomical positions from PCW12, we hypothesised that cell signalling mechanisms involved in generating motor activity (ion channels, calcium signalling, etc.) may not yet be fully matured. To this end, we tested whether the expression levels of ANO1, a calcium-activated chloride channel expressed in ICC and a major signalling mechanism vital to the establishment of pacemaker activity^15^, were delayed at earlier timepoints - even though key cell surface proteins (SCFR+) showed high expression and the intestinal segments were able to generate ongoing contractions. Interestingly, at PCW12, ANO1 staining was colocalised with SCFR+ in ICC (**Fig. 3 G, H, I**, arrows), ruling out any delayed expression of ANO1 in ICC to explain the different motor patterns in less developed human fetal small intestine. While the overall area of ANO1-stained structures in the gut wall did not change across time, the network-like appearance of ICC in the myenteric region (**Fig. 3 G** compared to **H & I**, arrows) appeared to mature between PCW12 and PCW16, suggesting that the interconnectivity of ICC and/or other signalling pathways may play a crucial role in establishing mature motor patterns in the developing intestine.

**Figure 3:**
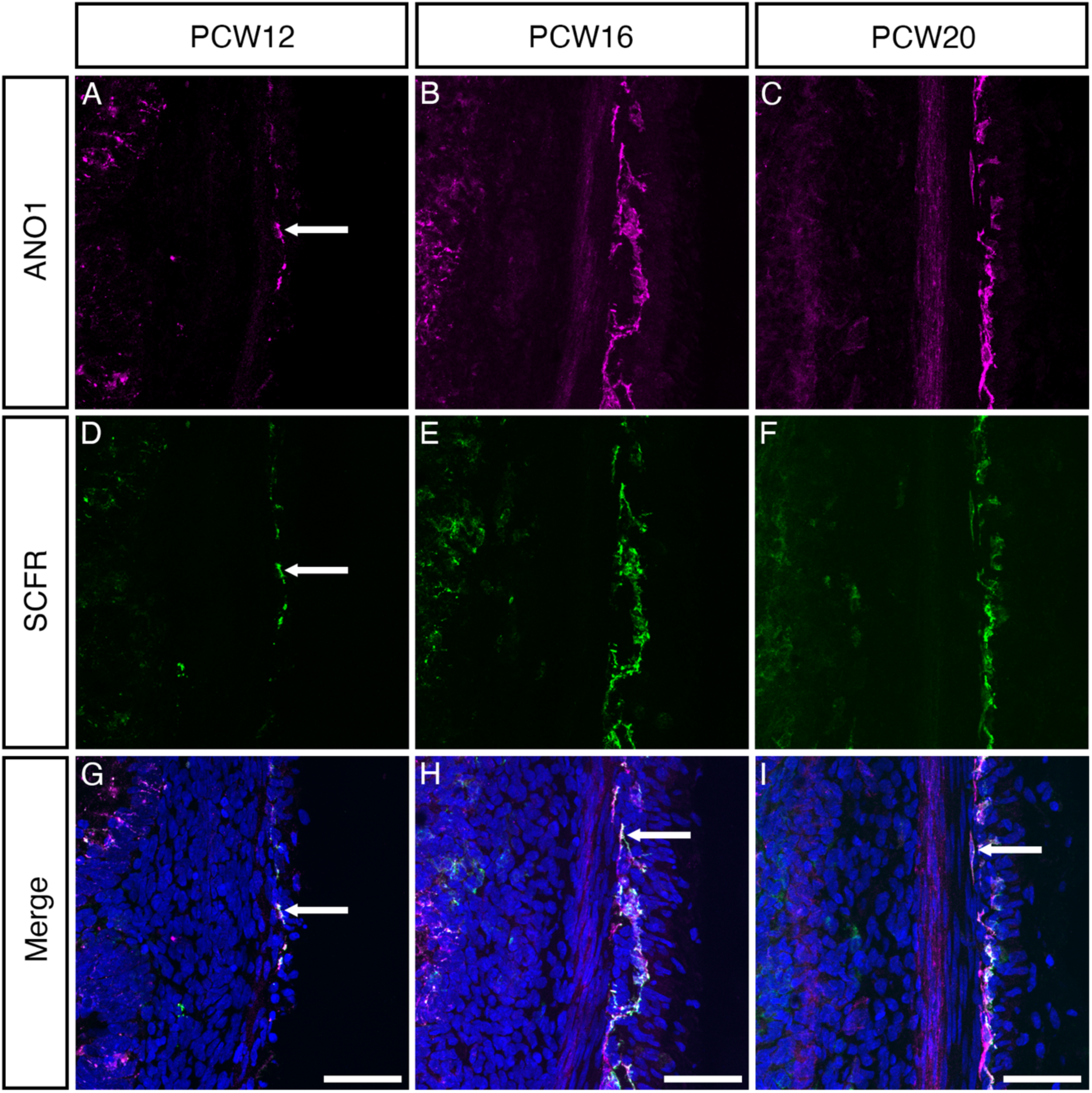
SCFR+ cells within the developing small intestinal wall express the calcium channel ANO1. (A-I) Representative confocal images of intestinal cryosections at PCW12, PCW16 and PCW20. ANO1 (magenta; A-C, G-I) and SCFR (green; D-I expression was observed small intestinal tissue at PCW12, PCW16 and PCW20. At PCW12, PCW16 and PCW20 the vast majority of ANO1+ cells colocalised for SCFR (G-I, arrows). Scale bars 50µM.

Hence, we then widened our search of genes that showed significant changes between PCW12 and PCW16/PCW20 to investigate other potential influences on developing motor patterns.

### Increasing contractile coordination in the human small intestine correlates with upregulation of the calcium and purinergic signalling pathways

To establish the potential molecular mechanisms which may contribute to the development of coordinated contractile patterns, in an unbiased and higher throughput fashion, bulk RNAseq analysis was conducted on n=3 samples for each age (PCW12, PCW16 and PCW20). Exploratory data analysis using unsupervised Principal Component Analysis (PCA) revealed that the samples clustered according to age (**Fig. 4A**), with PCW16 and PCW20 clustering more closely to each other than to PCW12. By adopting a generalised linear model, we could identify genes that differed between any of the developmental conditions in a single step. This approach revealed a total of 2019 differentially expressed genes (DEGs) with an FDR<0.05 **(Supplementary data file 1)**. We then clustered all DEGs based on their pattern of gene expression changes across the three timepoints. This analysis identified 5 unique expression patterns (**Fig. 4B**, cluster 1: 391 genes; cluster 2: 832 genes; cluster 3: 402 genes; cluster 4: 182 genes; cluster 5: 212 genes). Genes within each cluster were subjected to over-representation analysis to identify enriched processes or pathways using the Gene Ontology^16^ and KEGG (Kyoto encyclopaedia of genes and genomes^17^) databases (**Supplementary data file 2)**. We focussed our attention on cluster 2 (**Fig. 4B**) as this cluster contained >two-fold more genes when compared to other clusters. Within cluster 2, we found multiple enriched pathways relating to development of coordinated contractile activity, including calcium signalling pathways, the cAMP signalling pathway, the PI3K-Akt signalling pathway and the cholinergic synapse signalling pathway, in addition to pathways involved in vascular smooth muscle contraction and gastrointestinal smooth muscle contraction (**Fig. 4C**). Of particular relevance, the cholinergic synapse (**Fig. 4D**), gastrointestinal smooth muscle contraction (**Fig. 4E**) and vascular smooth muscle contraction pathways (**Fig. 4F**) all showed substantial upregulation of numerous key genes at PCW16 and PCW20 compared to PCW12.

**Figure 4:**
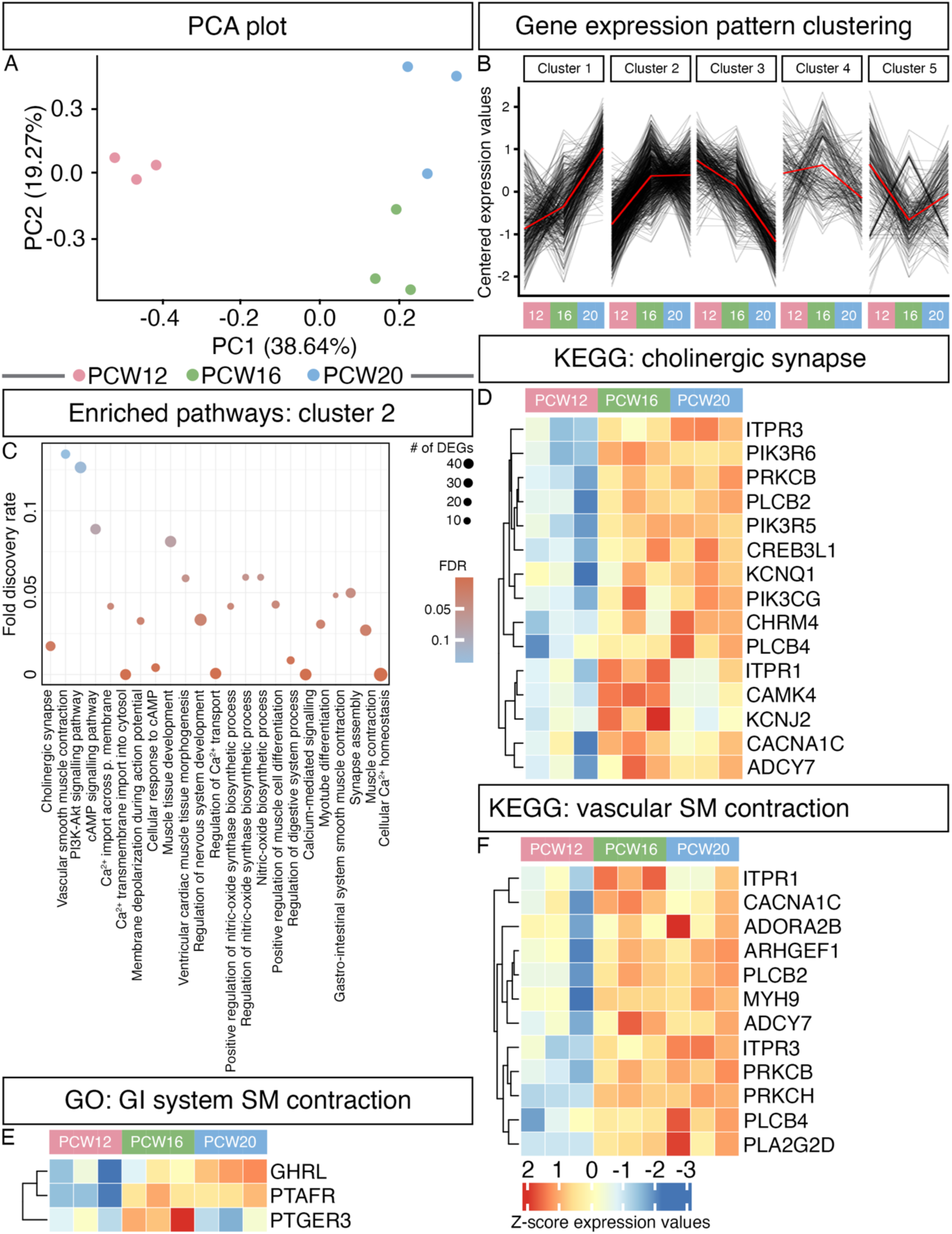
Gene expression pattern analysis of human small intestinal tissue reveals dramatic changes between PCW12 and PCW16/PCW20. (A) Unsupervised Principal Component Analysis (PCA) revealed that the samples clustered according to age (n=3 samples for each time point). (B) Differentially expressed genes between PCW12, PCW16 and PCW20 small intestinal tissue were clustered according to their expression profile, revealing 5 distinct expression patterns (cluster 1: 391 genes; cluster 2: 832 genes; cluster 3: 402 genes; cluster 4: 182 genes; cluster 5: 212 genes). (C) Over representation analysis of cluster 2 using the gene ontology and KEGG databases revealed enrichment of multiple biological pathways of interest, including muscle tissue development, regulation of calcium ion transport and positive regulation of nitric oxide synthase biosynthetic process, among others. (D-F) Heat maps showing the temporal expression pattern of differentially expressed genes in candidate KEGG and GO pathways at PCW12, PCW16 and PCW20 (n=3 samples for each time point), including cholinergic synapse (D), gastro-intestinal system smooth muscle contraction (E) and vascular smooth muscle contraction (F).

A preliminary investigation of the 2019 DEGs revealed a trend towards increased expression of genes associated with ion channels at PCW16 and PCW20 compared to PCW12 (**Fig. 5A**). Similarly, we detected increased expression of genes associated with ENS development at PCW16 and PCW20 compared to PCW12 (**Fig. 5B**). However, no obvious trends were found within DEGs associated with gap junctions or nNOS signalling (**Fig. 5C and D**, respectively).

**Figure 5:**
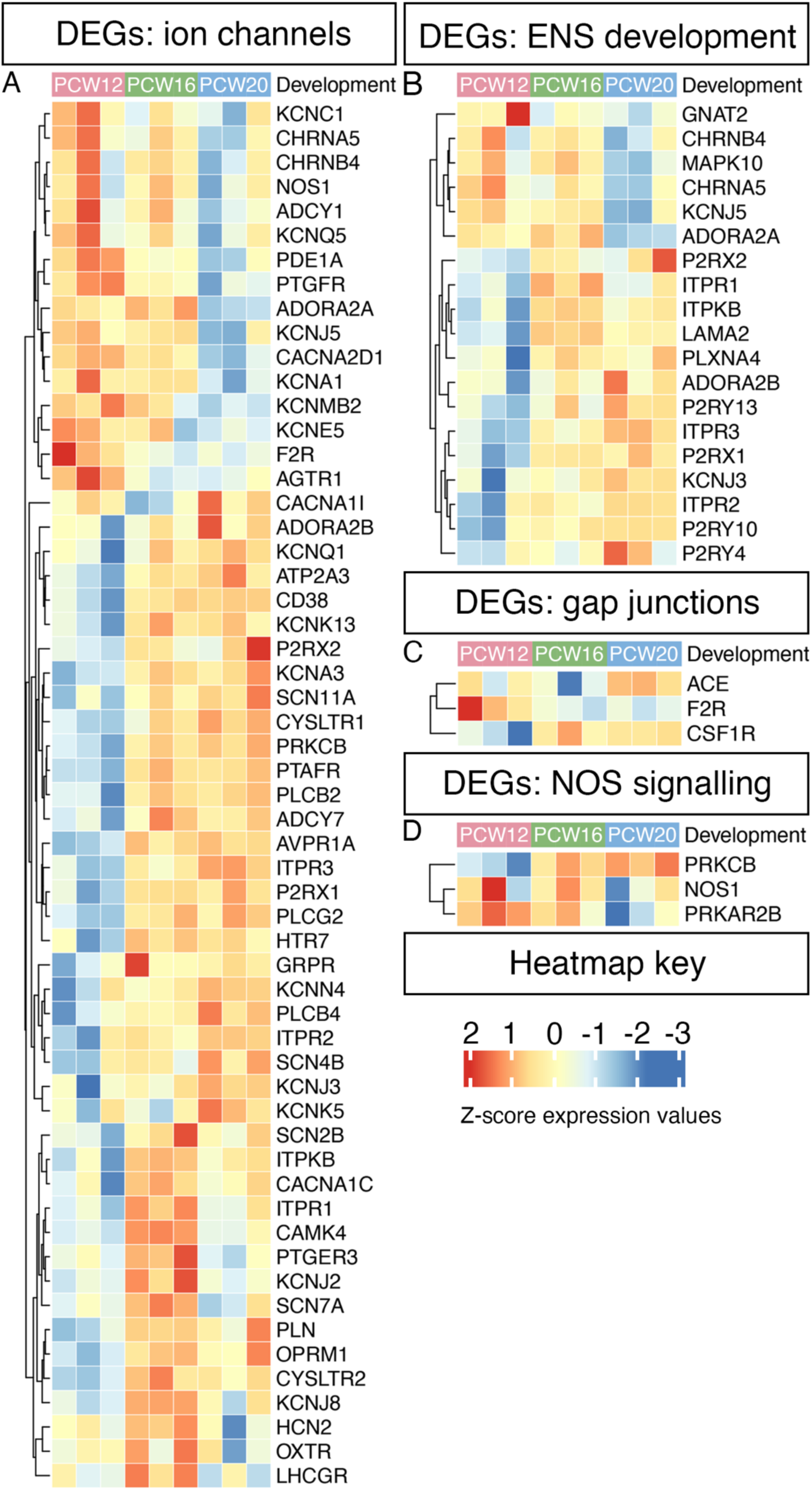
Analysis of differentially expressed genes highlights a correlating increase in genes associated with ion channels and enteric nervous system development with increasing age. (A-D) Heatmap representation of differentially expressed genes involved in ENS development and function. A generalised linear model revealed a trend towards an increased expression of ion channels genes (A) and genes associated with enteric nervous system development (B) at PCW16 and PCW20 tissue, compared to PCW12 tissue. However, no obvious trend was observed in the expression of genes involved in gap junctions (C) or nNOS signalling (D)

Upon further inspection it became clear that many of the significantly upregulated genes identified by this analysis were involved in the purinergic signalling pathway. Based on these observations, we performed gene set enrichment analysis^18^ on a selected subset of pathways (**Fig. 6A**) to assess their significance (a complete list of all pathways investigated can be found in **Supplementary data file 3**). Of all pathways examined, only the calcium signalling pathway, at PCW16 (normalised enrichment score 1.47, FDR=0.038) and the purinergic signalling pathway at PCW20 (normalised enrichment score 1.50, FDR=0.14) were observed to be upregulated when compared to PCW12, respectively (**Fig. 6A-C**).

**Figure 6:**
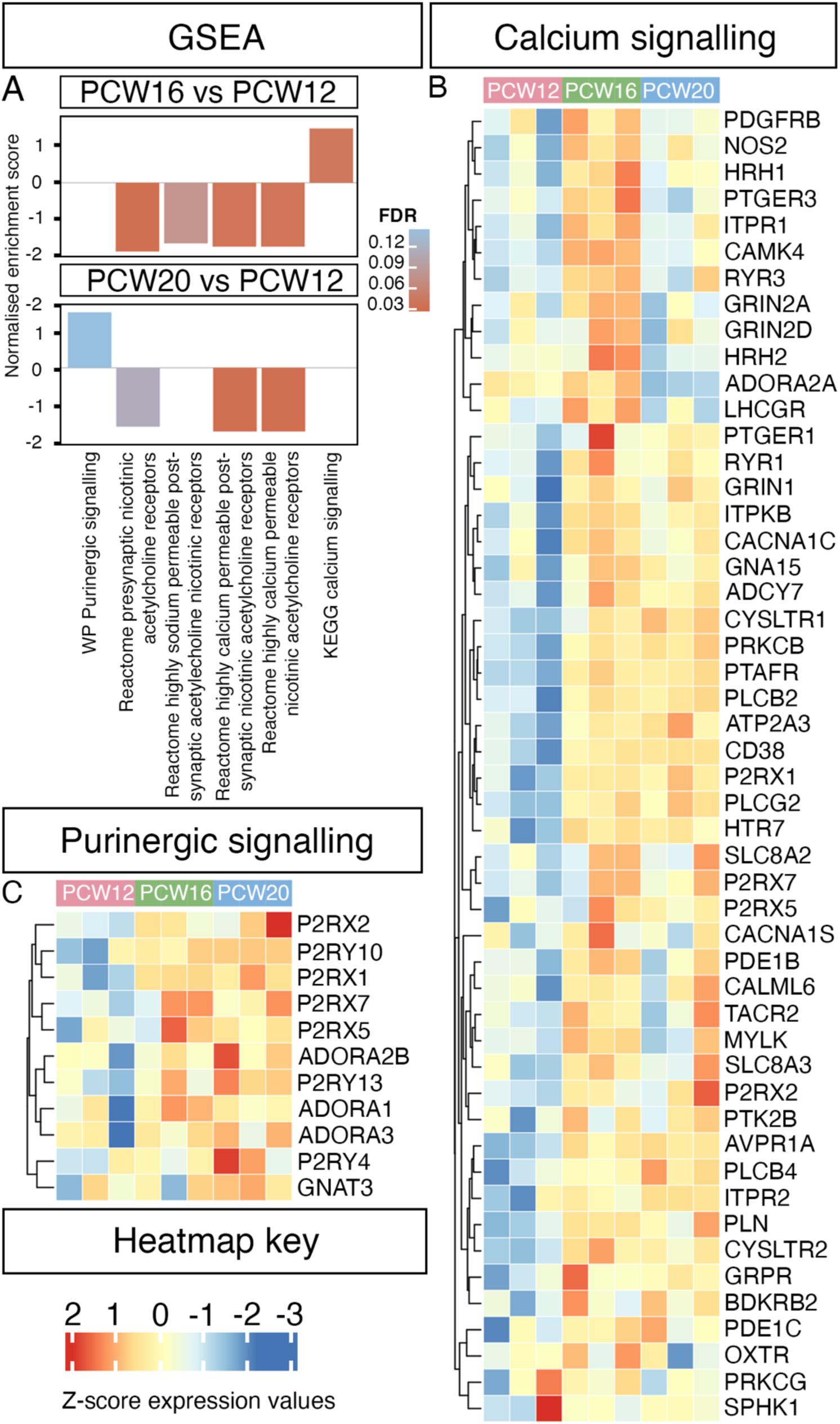
Expression of purinergic and calcium signalling pathways become significantly upregulated between PCW16 and PCW20. (A) Gene set enrichment analysis of specific candidate pathways revealed a significant upregulation of genes involved in calcium signalling at PCW16 compared to PCW12. At PCW20, expression of genes involved in purinergic signalling were significantly increased compared to PCW12. (B-C) Heatmaps of the leading-edge genes for both calcium signalling (B) and purinergic signalling (C) revealed a strong trend of increasing expression of genes with increasing age of human small intestinal tissue.

Calcium signalling is vital for numerous physiological functions, including smooth muscle contraction. Within the calcium signalling pathway gene set we identified several significantly upregulated genes of particular interest (**Supplementary data table 1**), including ITPR1 and ITPR2. ITPR1 and ITPR2 are inositol 1,4,5-triphosphate (IP_3_) receptors enabling release of intracellular calcium vital for smooth muscle contraction. Concordantly, we also detected upregulation of phospholipase enzymes required for the generation of IP_3_: PLCB2 and PLCB4. CACNA1C, the alpha subunit of the L-type voltage-sensitive Ca2+ channel that facilitates increased influx of calcium into a cell following membrane depolarization, was also increased. The ryanodine receptors RYR1 and RYR3 were upregulated, potentially facilitating release of intracellular calcium from the sarcoplasmic reticulum, as was the calcium-activated protein kinase PRKCB. Within the calcium signalling pathway we also detected several purinergic genes, including the fast responsive ion channel purinergic receptors P2RX1 and P2RX7, as well as the adenosine receptor ADORA2A.

Purinergic signalling from the ENS to SMCs, mediated by interstitial cells including ICC and PDGFRα+ cells, has been known to play a vital role in GI physiology since the 1960s^19^. We identified several genes of interest within the significantly upregulated purinergic signalling gene set (**Supplementary data table 1**), including both fast responsive ion channel purinergic receptors such as P2RX1, P2RX2 and P2RX7 as well as the slower responding, g-protein coupled purinergic receptors P2RY13, P2RY10 and P2RY4. We also detected upregulation of the adenosine receptors ADORA2B and ADORA3.

### Protein expression of key genes in the calcium and purinergic pathways is upregulated between PCW12 and PCW20 in the human small intestine

To determine whether the age-related changes in RNA expression could also be detected with spatial resolution at the protein level, quantitative immunofluorescent analysis was performed on human small intestine at the same ages (PCW12, PCW16 and PCW20, **Fig. 7**). Following a thorough literature review, four targets were chosen from the lists of genes within the calcium and purinergic signalling pathways for further expression analysis based upon potential key roles in the development of contractile coordination: P2RX2 and ADORA2B (both involved in the purinergic pathway) and ITPR1 and CACNA1C (both involved in the calcium signalling pathway). Immunofluorescent investigations were conducted on cryosectioned tissue (n=3 for each target). We observed a significant increase in P2RX2 expression between PCW12 and PCW20 (PCW12: 20.27 ±12.62%, PCW16: 73.72 ±16.64%, PCW20: 105.8 =/-19.88%, p=0.0254, **Fig. 7A-D**). At PCW12 and PCW16, P2RX2 appeared restricted to the presumptive myenteric plexus region (**Fig. 7A&B,** arrows) whereas at PCW20 expression was observed both within the myenteric and submucosal plexus regions in presumptive ganglia-like clusters (**Fig. 7C**, arrowheads). ADORA2B expression was significantly increased at PCW20 (125.1 ±11.89%) compared to both PCW12 (12.45 ±5.704%, p=0.0003) and PCW16 (15.18 ±9.134%, p=0.0004, **Fig. 7E-H**) with expression at PCW20 again appearing localised to the myenteric and submucosal plexus regions (**Fig. 7G**, arrowheads) though at a lower level than P2RX2. ITPR1 expression increased progressively across all three time points under examination, with a significant increase between PCW12 and PCW20 (PCW12: 34.12 ±15.41%, PCW16: 101.7 ±31.07%, PCW20: 125.8 ±9.683%, p=0.0470, **Fig. 7I-L**). At PCW12, sporadic labelling of ITPR1 was observed within the submucosal region (**Fig. I,** arrowheads) and at the serosal surface of the gut (**Fig. I,** arrows). At PCW16, more robust ITPR1 expression was observed which appeared to be restricted within the myenteric and submucosal plexus regions in presumptive ganglia-like clusters (**Fig. 7J**, arrowheads). However, by PCW20 ITPR1 expression was observed almost ubiquitously throughout the submucosa and *tunica muscularis* (**Fig. 7K**). CACNA1C expression was significantly increased at PCW20 (23.86 ±6.218%) relative to both PCW12 (1.213 ±0.9972%, p=0.0124) and PCW16 (3.215 ±1.570%, p=0.0188, **Fig. 7M-P**). At PCW12, very little expression of CACNA1C was identified (**Fig. 7M**). By contrast, at PCW16 CACNA1C expression could be identified and was found to be largely restricted to the presumptive myenteric plexus region (**Fig. 7N**, arrows). With further maturation to PCW20 CACNA1C expression could more readily be identified, however, surprisingly this expression appeared to be more robust within the myenteric and submucosal plexus regions (**Fig. 7O**, arrowheads) with punctate staining observed in the smooth muscle regions (**Fig. 7O**, hollow arrows).

**Figure 7:**
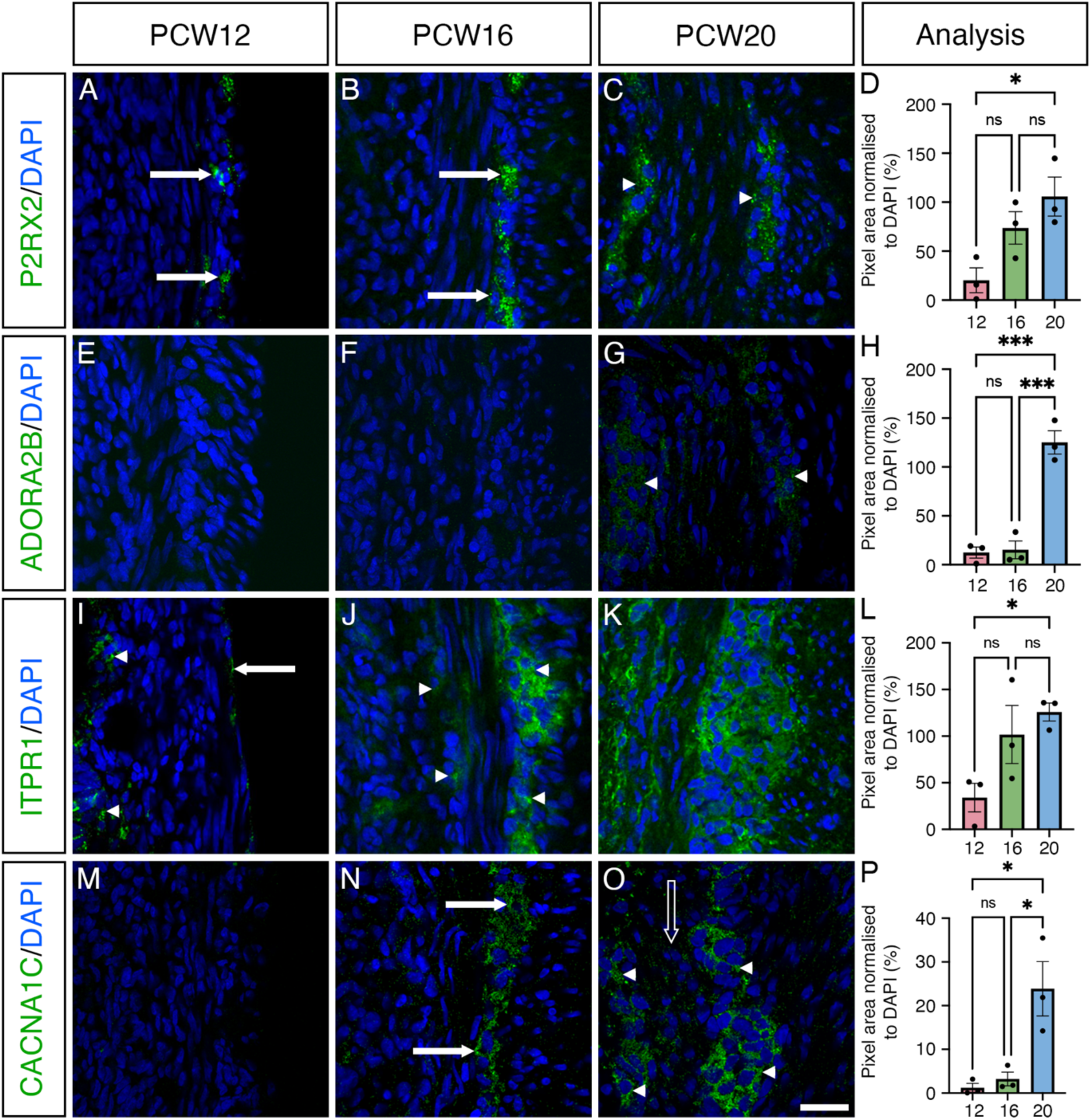
Expression of key proteins involved in the purinergic (P2RX2 and ADORA2B) and calcium (ITPR1 and CACNA1C) pathways are significantly increased at PCW20 compared to PCW12. (A-C) Representative fluorescent confocal images showing P2RX2 (green) and DAPI (blue) expression in cryosections of human small intestine at PCW12, PCW16 and PCW20. (D) Summary data showing P2RX2+ pixels expressed as a percentage of total DAPI staining (n=3 samples per age). (E-G) Representative fluorescent images showing ADORA2B (green) and DAPI (blue) across all timepoints. (H) Summary data showing ADORA2B+ pixels expressed as a percentage of total DAPI staining. (I-K) Representative fluorescent images showing ITPR1 (green) and DAPI (blue) across all timepoints. (L) Summary data showing ITPR1+ pixels expressed as a percentage of total DAPI staining. (M-O) Representative fluorescent images showing CACNA1C (green) and DAPI (blue) across all timepoints. (P) Summary data showing CACNA1C + pixels expressed as a percentage of total DAPI staining. Scale bars 25µM.

Taken together, transcriptional and protein expression analyses suggest that tight control of purinergic signalling and regulation of calcium dynamics are key factors in the establishment of coordinated motor patterns in the developing human gut.

## Discussion

The development of the full repertoire of motor behaviours in the small intestine is a tightly controlled process involving specific temporal expression of genes in parallel with morphological complexity as the gut develops. In this study, we highlight (1) the early-mid second trimester (PCW12-20) as a key window during which motility patterns undergo dramatic developmental shifts and (2) maturation of cell signalling pathways as a key mechanism underlying these changes. Importantly, our results suggest that between PCW12 and PCW20, as the musculature and ENS mature, inhibitory mechanisms exert an increasing influence to permit long propagating contractions. Our data suggest this adaptation involves purinergic signalling.

We show that human fetal small intestinal segments display contractile activity from as early as PCW12, with ongoing rhythmic propagating contractions that likely mix and propel intestinal contents. However, the dynamics of these contractions alter rapidly, with a dramatic increase in both the frequency and velocity of propagating contractions between PCW12 and PCW16. This increase may be at least partially driven by the fetus swallowing amniotic fluid. Gastric peristalsis appears and develops from 14-23 weeks of gestation, propelling ingested amniotic fluid into the intestines^12, 20, 21^. The observed increase in motor capabilities of the small intestine between PCW12 and PCW16 suggests the development of the small intestine may coincide with the greater need to clear ingested amniotic fluid from the GI tract.

The fetal motility we observed displayed regular, ongoing, slowly propagating small amplitude contractions, suggesting a strong myogenic basis. Notably, it was indistinguishable to motility patterns observed in rodent models following neutralization of enteric neural activity via addition of tetrodotoxin^22^. However, the consistency, regularity and coherence of the propagating contractions are indicative of excitation spread via ICC networks, rather than the haphazard timing and irregular spread of muscle action potential-mediated contractions observed in smooth muscle layers of tissue devoid of ICC^23^. Indeed, we detected both ICC and PDGFRα+ cells from PCW12, as well as ANO1, a key ion channel involved in ICC pacemaker activity. Of note, we were unable to detect differences in the overall area of ANO1-stained structures within the developing gut wall. However, the quality of the antibody staining in cryosections likely contributed to our failure to detect changes, given the fine filamentous projections of networked ICC. Interestingly, SI motility changed dramatically, with substantial increases in contractile frequency and velocity between PCW12 and PCW16. This correlated with robust upregulation of genes involved in ENS development between these timepoints, including specific pathways such as the cholinergic synapse pathway, suggesting that between these time points excitability within the ENS is maturing. This is in line with previous detection of compound ENS activity in the human fetal colon at PCW16^14^. However, we detected no further increase in contractile frequency and velocity between PCW16 and PCW20.

In murine studies, several processes have been found to contribute to regulation of GI peristalsis. Previous reports have shown the importance of prostaglandin signalling, via PGE2-PGER3 receptor activation, in slow wave propagation^24^. In line with this, our RNAseq analysis showed significantly increased expression of *PTGER3* at PCW16 compared to PCW12 (log fold change 0.62, p= 0.00008). However, we did not detect any significant change in expression of enzymes involved in PGE2 synthesis in the ages under examination. We also examined the potential role of other chronotropic agents found to have a role in gastric peristalsis, such as cholinesterases^25^, but the expression of key enzymes in the pathway of cholinesterases (such as *ACHE* and *BCHE*) also failed to show a significant alteration in the ages examined. Indeed, the observed alterations in SI motility correlated with significantly increased expression (by GSEA) of only two pathways: the calcium signalling pathway at PCW16 and purinergic signalling pathway at PCW20, which were observed below the false discovery rate, when compared to PCW12. The upregulation of these pathways may indicate an increasing influence of pacemaker-driven activity via the SIP-syncytium.

Calcium transients are a key feature of ICC-mediated pacemaker activity. We focussed on two genes, both of which appeared as a “hit” in multiple enriched pathways and are directly involved in ICC-SMC contraction coupling: ITPR1 and CACNA1C. ITPR1 encodes a ligand-gated ion channel facilitating calcium release from the endoplasmic reticulum^26^ and is known to be expressed in ICC^27^, with a crucial role in generating rhythmic Ca^2+^ transients that activate Ca^2+^-activated Cl^-^ currents to depolarize the ICC itself and any coupled cells. CACNA1C encodes an L-type voltage dependent calcium channel known to be expressed in SMCs^28^. Disruption of CACNA1C activity in mice has been shown to lead to impaired GI contractility, possibly via uncoupling SMC-ICC^29^. The importance of these genes in GI motility is reinforced by the severe GI motility defects observed in patients with ITPR1^30^ or CACNA1C^31^ mutations.

While purinergic signalling within the ENS, such as neuron-neuron transmission during the peristaltic reflex and migrating motor complexes (MMCs), has been well characterised, synaptic transmission from the ENS to SMCs is mediated by interstitial cells^32^. We observed increased expression of numerous purinergic signalling-related proteins in PCW20 tissue compared to PCW12 tissue, including P2RX1, P2RX2, P2RX7, P2RY4, P2RY13 and ADORA3, all of which have been previously shown to be expressed by murine ICC^33^. This increased expression likely reflects the increasing capability of ICC to respond to purinergic synaptic inputs during later development. Notably, P2RX2 expression has been identified in ICCs in the guinea pig and mouse ileum, where it was hypothesised to contribute to pacemaking^34^, supporting our findings. Purinergic inhibition is an important component of both tonic inhibitory and descending inhibitory pathways that relax the gut musculature –however, this also prevent spurious “ectopic” contraction initiation sites from forming that interfere with the propagation of migrating motor complexes or slow-wave mediated contractions. Our finding that the number of initiation sites dramatically decreases at PCW20 correlates well with our observations noting prominent increases in purinergic proteins at the same age.

In addition to ICC, PDGFRα+ cells are a crucial component of ENS-driven purinergic signalling, playing a critical role in tonic inhibition of GI contractile excitability^35^. Excitatory and inhibitory enteric motor neurons of the ENS synapse both directly onto SMCs as well as indirectly via interstitial cells^6^, with neuronally-driven purinergic signalling causing hyperpolarization of PDGFRα+ cells and inhibiting SMC contractions^35, 36^. The observed increase in purinergic signalling pathway expression may act to prevent aberrant contractility. Indeed, ADORA2B, one of the genes highlighted as upregulated at PCW20 compared to PCW12 in our study, appears to play a role in mediating contractility^37^. ADORA2B encodes a G-protein coupled receptor expressed by both epithelial cells and SMCs in the GI tract^33^ and *Adora2b*^-/-^ mouse models display impaired colonic relaxation^38^. These results highlight the important role of interstitial cells, and the inhibitory capabilities of the ENS, in the development of coordinated GI motility during the second trimester and may help to identify pharmacologic modulators of gastrointestinal motility disorders. Further, the dramatic reduction in initiation sites we observed at PCW20, compared to earlier timepoints, likely impacts on the propagation of pacemaker activity. Pacemaker activity spreads in an isotropic fashion (radiating outwards at a constant velocity from the site of initiation). Therefore, the primary determinate of whether contractions propagate in the oral or anal direction is the location of the initiation site. Hence, it appears that the key to maintaining coherent propagating contractions over long distances is to prevent “ectopic” pacemaker initiation sites from forming elsewhere along the segment. Such sites can form due to wall stretch, mucosal stimulation, neural excitation and in response to inflammatory mediators^24^. Again, this could be at least partially explained by our observed increase in purinergic signalling at PCW20 compared to PCW12. The maturation of this descending inhibitory pathway may facilitate tonic inhibition and relaxation during reflexes and complexes anal to the point of stimulation, helping to ensure that propagating contractions are not impeded by ectopic pacing sites and thereby can travel long distances away from their site of initiation^39, 40^.

This study sought to determine the timeline and molecular mechanisms underlying the development of coordinated contractility in the human fetal intestine. The sourcing of appropriate tissue, at the correct developmental timepoints, for functional assessments is notoriously difficult in human studies. Nevertheless, as opposed to alternative studies reporting activity with low numbers (e.g., n=1 sample/fetal age) we have provided data across multiple (n≥3) replicates for each developmental stage and analysis performed. Of note, the consistency of the contractile patterns observed across multiple fetal intestinal samples permits a high level of confidence in our findings. To assess the molecular mechanisms underlying the development of coordinated contractile activity we performed bulk RNAseq which provided a holistic representation of global changes in gene expression within the human fetal intestine. Importantly, such a bulk approach fails to provide the granularity of single-cell or spatial transcriptomic approaches which may have provided information on transcriptional expression within specific cell types, although such techniques are also not without limitations.

In conclusion, this work suggests that the human fetal small intestine is motile at PCW12 and undergoes dramatic alterations in the coordination of motor patterns by PCW16, with further maturation of coordinated activity between PCW16 and PCW20. These changes in motility occur after the major cell types involved in rhythmic GI motility (enteric neurons, ICC, PDGFRα+ cells and SMCs) have already differentiated and/or formed networks. Crucially, we show that these changes correlate with increased upregulation of the calcium and purinergic signalling pathways, highlighting the role of interstitial cells in developing GI motility at early stages in human development. Finally, we identify multiple molecular targets that appear to be key for the development of fetal intestinal motor patterns and could therefore be vulnerable to insult and downstream dysfunction. It is probable that even minor perturbations in GI development can lead to significant motility issues, as evidenced by the significant proportion of the global population suffering from motility complaints^1^. Hence, greater understanding of the developmental processes involved in establishing motor patterns in the human intestine will be vital to produce novel therapeutic options to treat/manage gut motility disorders.

## Supporting information

Manuscript (including Supplementary Figures)

## Author contributions

B.J., A.C., L.P., S.P., B.C. and P.T. acquired and interpreted data. B.J., P.V.B., G.W.H., and C.J.M. contributed to study concept and design and interpreted data. C.J.M. acquired funding. B.J., L.P., S.P., G.W.H., and C.J.M. drafted the manuscript.

## Data availability statement

All sequencing data have been deposited on NCBI’s SRA and at GEO and be made available upon publication

---

**Supplementary Figure 1:**
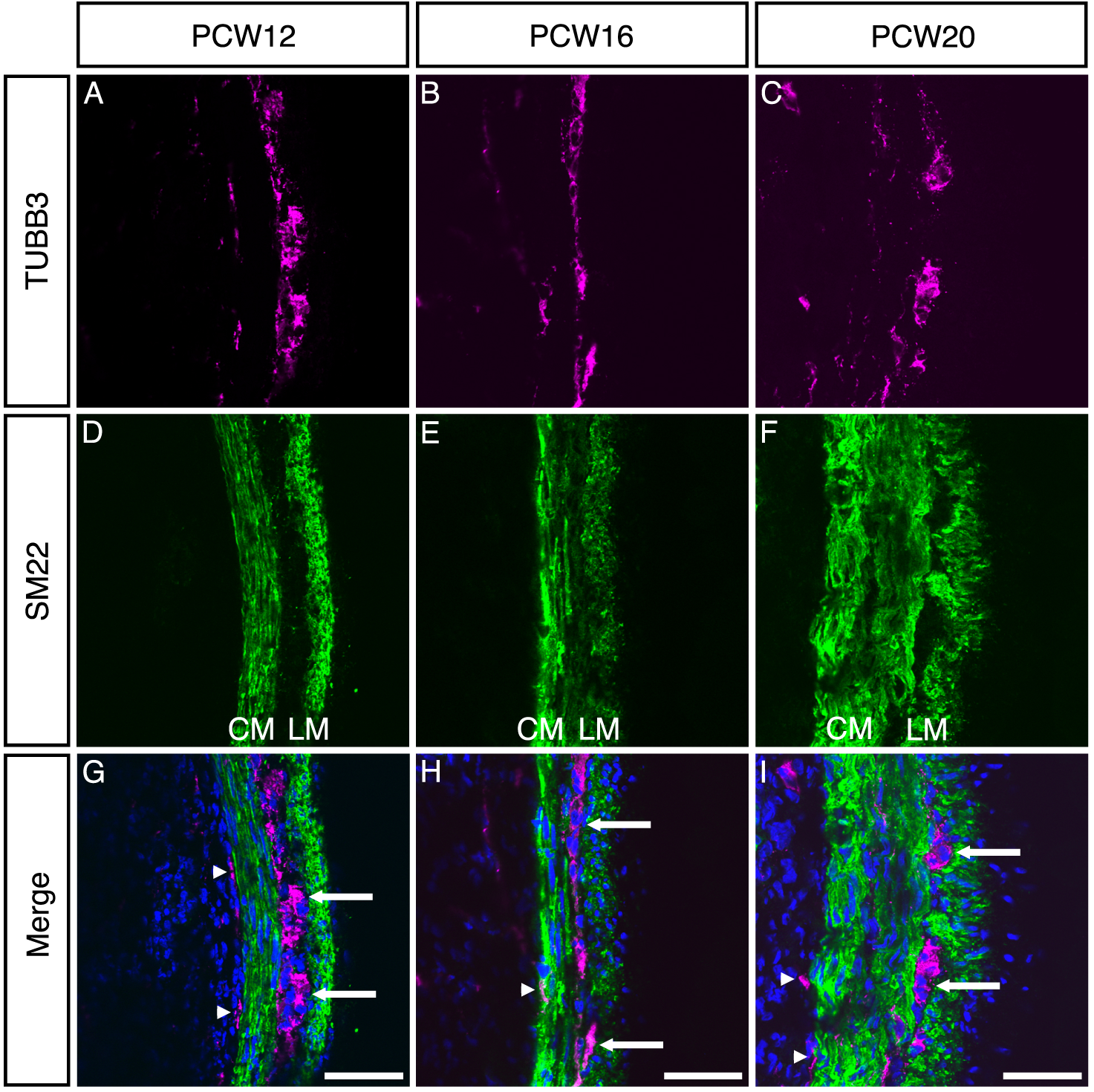
Neurons are present in the small intestinal wall from PCW12 in both the myenteric and submucosal plexuses and condense into obvious ganglia by PCW20. (A-I) Representative confocal images showing expression of TUBB3 (magenta), SM22 (green) and DAPI (blue) at PCW12, PCW16 and PCW20. At PCW12, TUBB3+ neurons could be clearly seen, organised in concentric rings within the small intestinal wall (A-C). Co-staining with the smooth muscle cell marker SM22 revealed the circular muscle layer (CM) and longitudinal muscle layer (LM) were also present at PCW12 (D-F). At PCW20, the muscle layers had thickened considerably (F). TUBB3+ neurons were clearly organised into the myenteric (arrows) and submucosal (arrowheads) plexuses (G-I). By PCW20, neurons, particularly of the myenteric plexus, had coalesced more obviously into ganglia (I, arrows). Scale bars: G-I = 50µM.

**Supplementary Figure 2:**
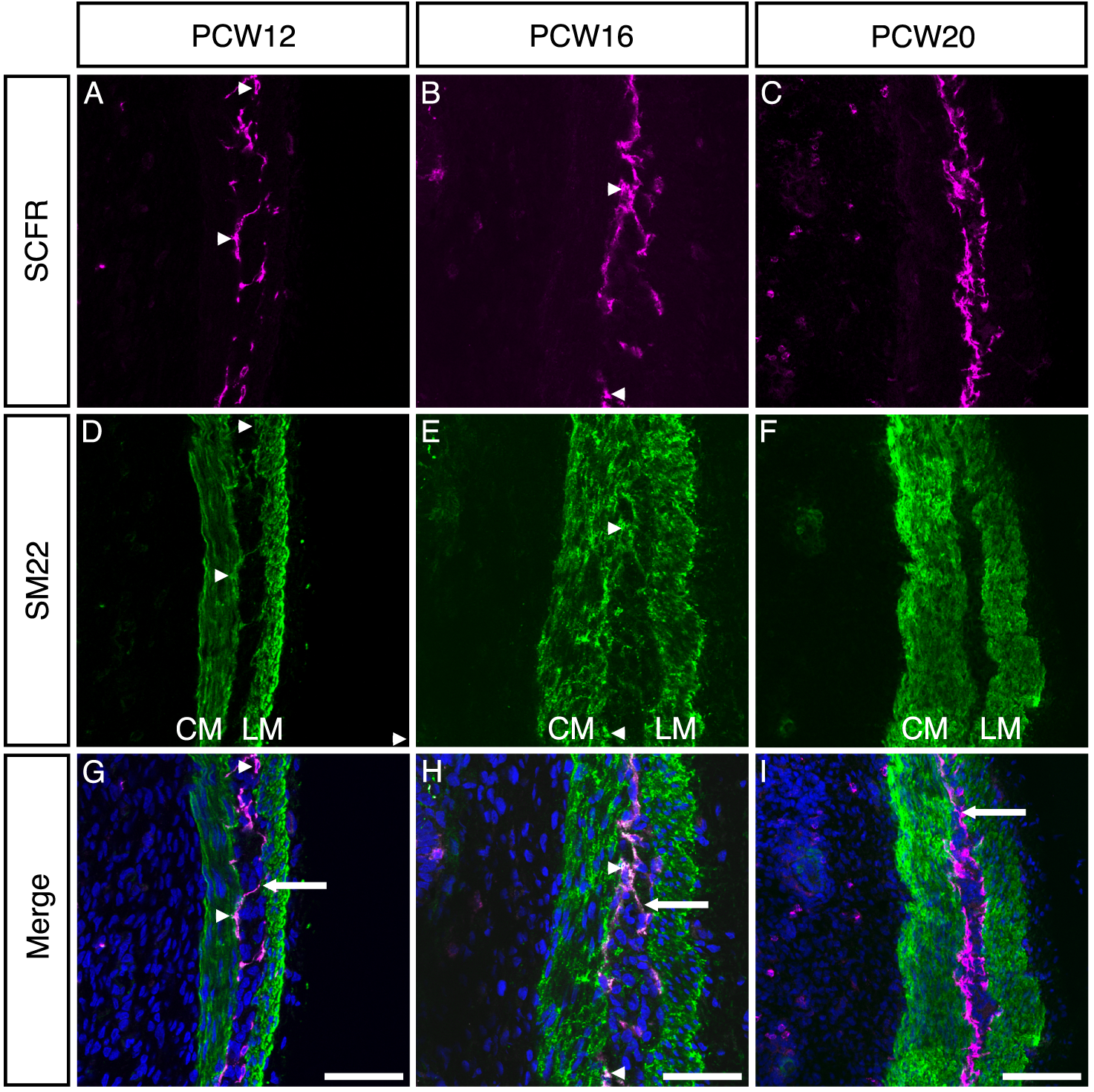
Interstitial Cells of Cajal are present in the small intestinal wall from PCW12 and appear to ‘line’ the inner separation of the circular and longitudinal muscle layers. (A-I) Representative confocal images showing expression of SCFR (magenta), SM22 (green) and DAPI (blue) at PCW12, PCW16 and PCW20. SCFR+ ICCs were present within the SM22 positive musculature at PCW12, PCW16 and PCW20 (A-C), located predominantly between the circular (CM) and longitudinal (LM) muscle layers (D-F). Often, the SCFR+ cells extended from one muscle layer to the other, forming ‘bridging’ connections (G-I, arrows). SCFR+ cells were also positive for SM22 (A, D, G, arrowheads). SCFR+ cells increased in density between PCW12 and PCW20 and appeared to ‘line’ the inner surfaces of the SM22 positive muscle between the circular and longitudinal muscle layers (G-I). Scale bars: G-I = 50µM.

**Supplementary Figure 3:**
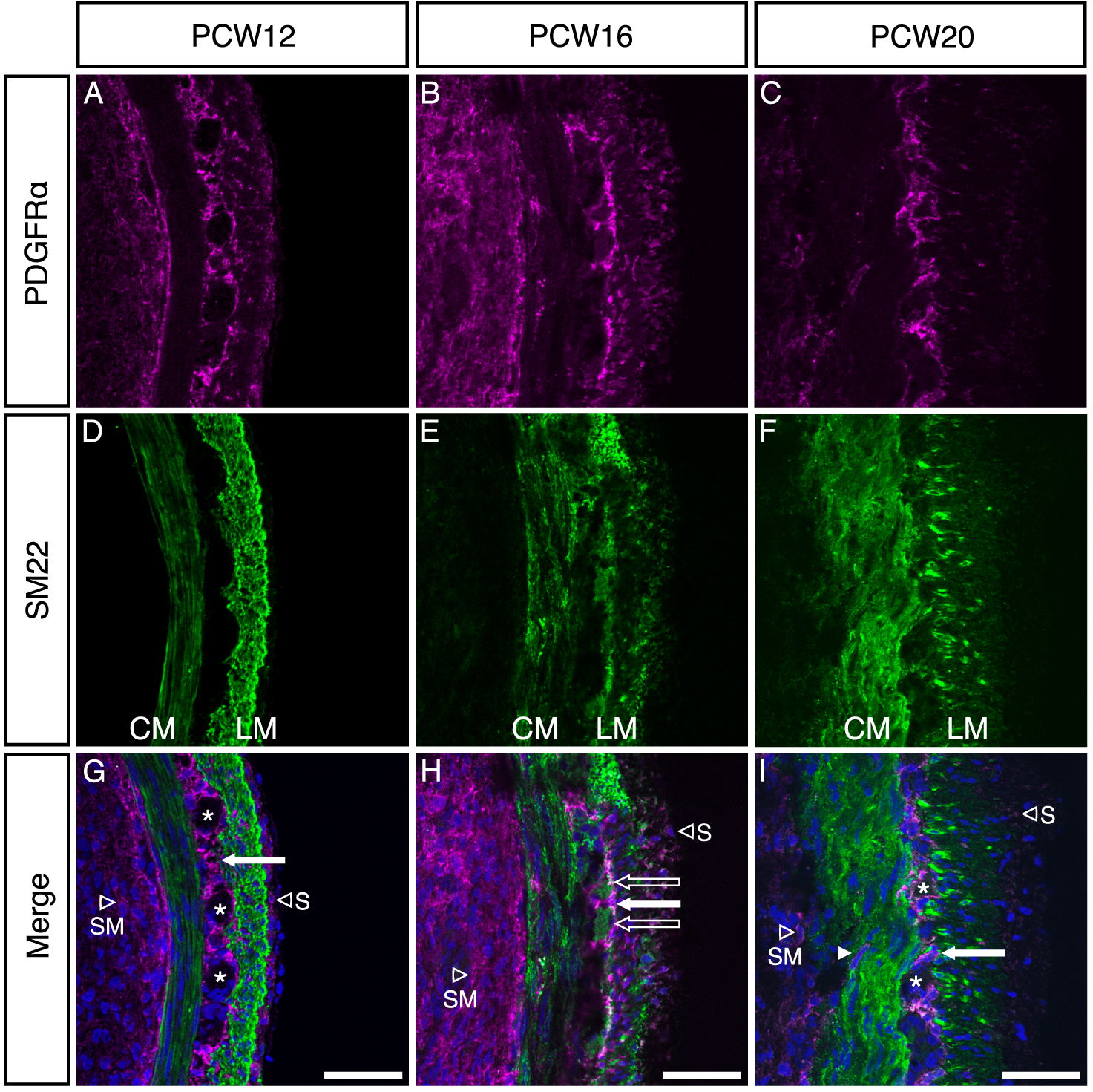
PDGFRα positive cells are present in the small intestinal wall from PCW12. (A-I) Representative confocal images showing expression of PDGFRα (magenta), SM22 (green) and DAPI (blue) at PCW12, PCW16 and PCW20. PDGFRα+ cells were observed in the small intestinal wall at PCW12, PCW16 and PCW20 (A-C). The majority of PDGFRα cells were found between the circular and longitudinal muscle layers (G-I, arrows), but were also found to a lesser degree in the submucosa (G-I, hollow ‘SM’ arrowheads) as well as on the serosal edge (hollow ‘S’ arrowheads). PDGFRα+ cells often appear to envelope other structures, including several which did not stain positive for SM22 (G, I, asterisks) as well as some SM22+ structures (H, hollow arrows). Scale bars: G-I = 50µM.

**Supplementary Figure 4:**
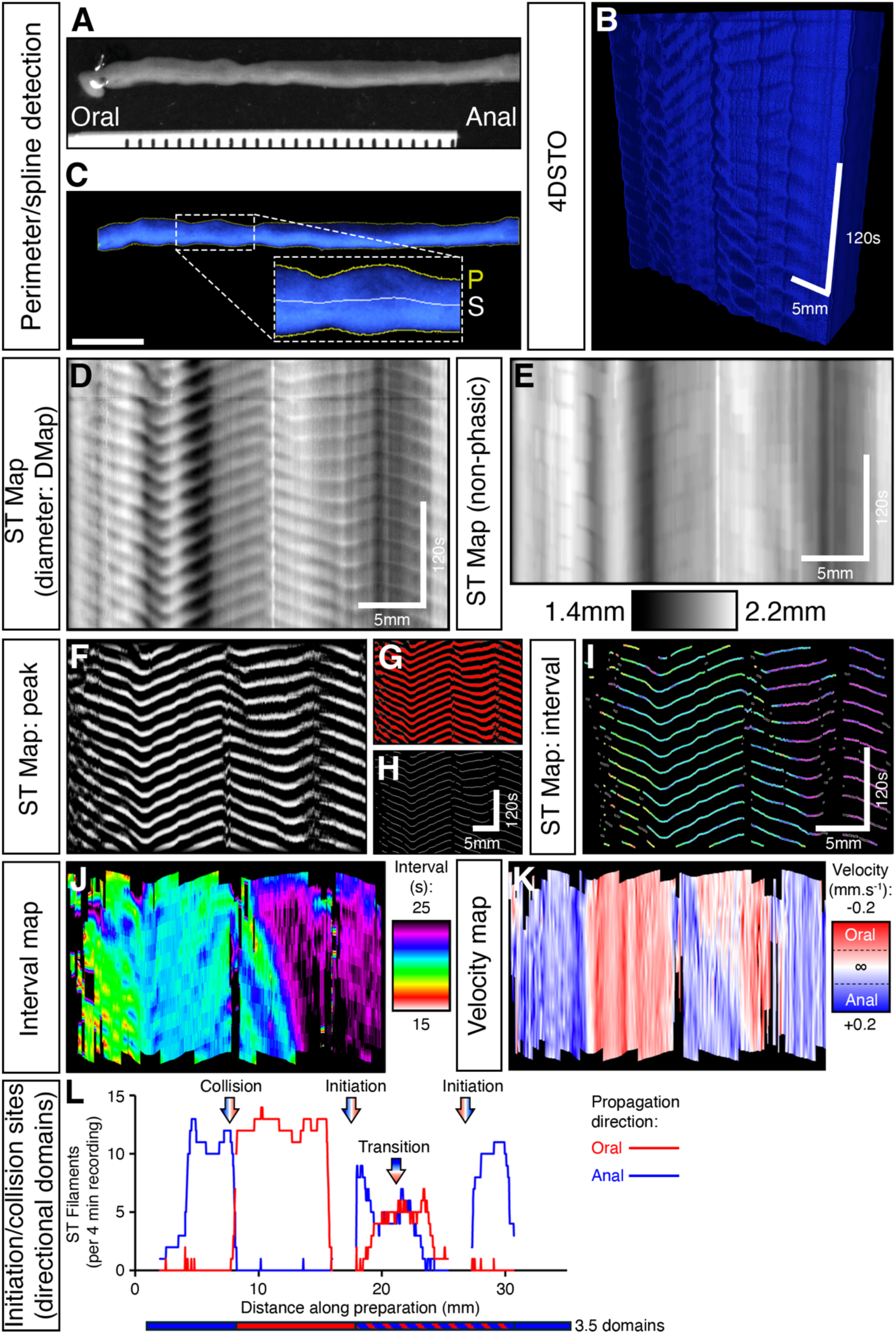
Image analysis techniques and workflow to extract motility characteristics from segments of isolated intestine. (A-L) Representative images showing the processing steps used to generate spatio-temportal maps and extract functional data. (A) Representative frame from a recording showing an isolated segment of small intestine aligned in the horizontal axis. Movies were calibrated using the ruler included in the field of view (FOV). After pre-processing and conversion to coordinate-based PTCLs (see Methods), a four-dimensional spatio-temporal object (4DSTO) was constructed (B). The outer perimeter of the intestinal segment was located, then a mid-line spline was added to guide the calculation of intestinal diameter (C). The diameter along the segment was determined for every slice (frame) and color-coded according to diameter (small diameter = black, large diameter = white). The resulting line of diameter from each slice were stacked underneath each other creating a ST Map (D) with the distance along the segment in the X-axis, time in the Y-axis and the color-coded diameter as the level of grayscale. Contractions are visualized as sloped darker lines. To aid the extraction of phasic contractions, the degree of background distension was calculated by locating diameter minima in a long time window (20s: see E). Following subtraction of non-phasic background distension, a recursive peak detection algorithm was applied to locate the points of maximum contraction or dilation (F). Following thresholding (G) and skeletonization (H), ST maps containing contraction trajectories were used to calculate the frequency of phasic contractions (wave-to-wave interval), with interval values color-coded as a spectrum (short interval = high frequency = white/red; long interval = low frequency = black/purple) (I). Linear interpolation between color-coded contraction trajectories was used to create FMaps (wave-to-wave interval) (J). Using a similar approach, calculation of the instantaneous slope along each contraction trajectory allowed the direction and velocity of each contraction as it propagated along the intestinal segment to be calculated and mapped (K). (L) The frequency of contractile initiations and collisions was determined for each sample. The red-blue shifts within the arrows indicates the transition of propagation direction. The duration of contraction propagation in each direction is summed in the ‘domains’ bar under the x axis.

**Supplementary Figure 5:**
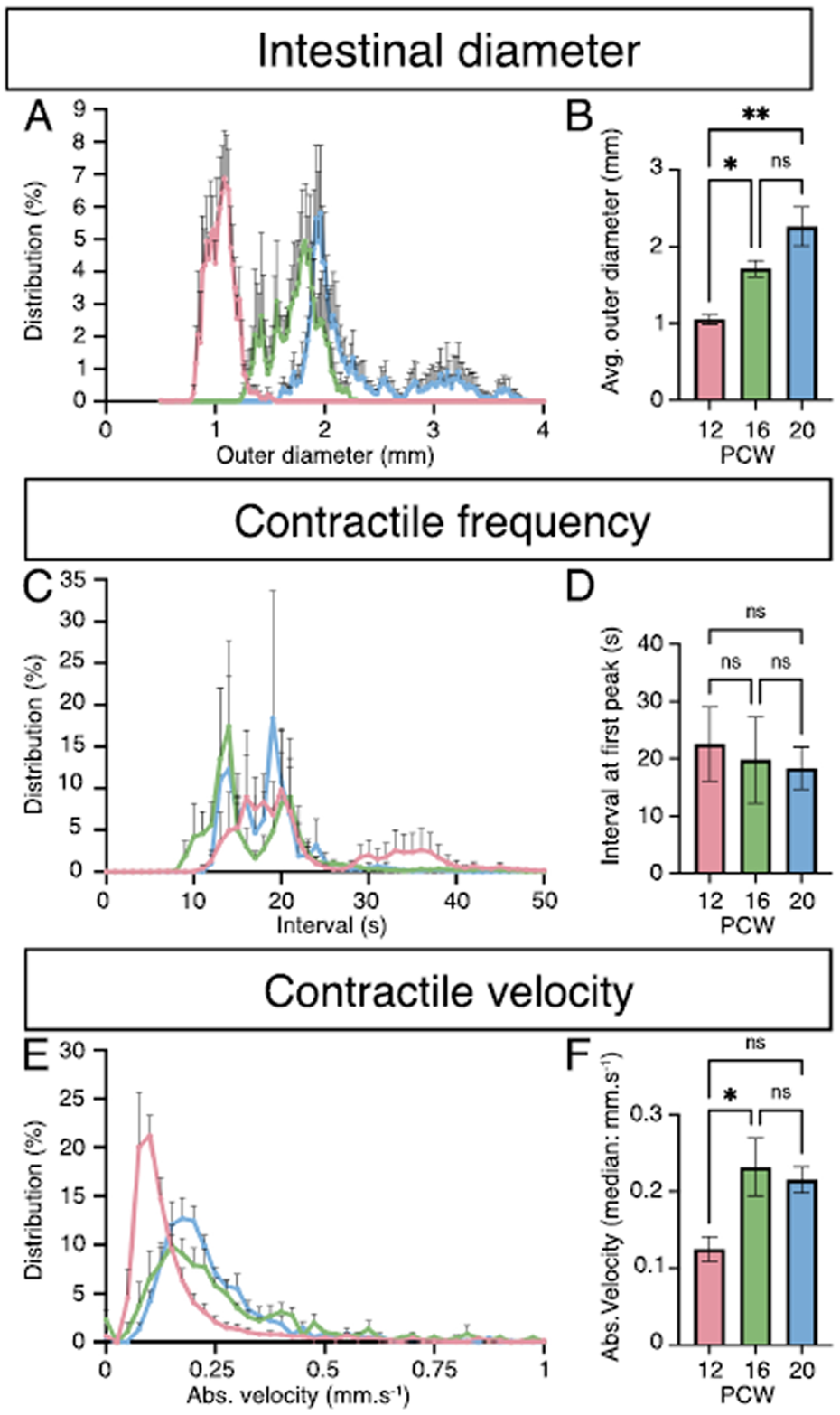
Summary statistics of contractile activity. (A-F) Summary data showing intestinal diameter (A-B), contractile frequency (C-D) and Contractile velocity (E-F) in PCW12, PCW16 and PCW human fetal intestine. (A-B) Average outer diameter values from PCW12, PCW16 and PCW20 intestinal segments (n=4 for each group). The outer diameter of intestinal segments doubled from PCW12 to PCW16/20 (*p<0.05, **p<0.01; B). (C-D) The wave-to-wave interval was shorter (faster frequency) in intestinal segments from PCW16/20 compared to PCW12. In particular, extremely long intervals between contractions were observed in intestinal preparations from PCW12 (see pink trace > 30s). Due to the variability in wave-to-wave interval due to the number and position of contraction initiation sites, these values did not reach statistical significance (E). (E-F) The velocity propagating contractions doubled from PCW12 to PCW16 (*p<0.05; F).

## Abbreviations

DEGs: Differentially expressed genes
ENS: Enteric nervous system
FDR: False discovery rate
GI: Gastrointestinal
GSEA: Gene set enrichment analysis
ICC: Interstitial cells of Cajal
KEGG: Kyoto encyclopaedia of genes and genomes
PFA: Paraformaldehyde
PDGFRα: Platelet-derived growth factor receptor alpha positive cells
PCW: Post-conception week
PCA: Principal Component Analysis
RT: Room temperature
SIP: **<u>S</u>**MCs, **I**CC, **<u>P</u>**DGFRα+ cells
ST Map: Spatiotemporal maps
SCFR+: Stem cell factor receptor positive

