## Supplementary material for "Purinergic signalling and calcium dynamics: potential drivers in the onset of coordinated intestinal motility in human fetal development": Manuscript (including Supplementary Figures)

**Supplementary Materials and Methods**

**Tables**

**Supplementary methods table 1: all human samples used in study. Note that some samples were used in multiple experiments (e.g., live imaging and immunohistochemical quantification).**

| All human samples used | | | | |
|---|---|---|---|---|
| **Sample code** | **Age** | **Karyotyping** | **Gender** | **Experiment** |
| 15316 | PCW12 | rsa(13,15,16,18,21,22,X)x2 | Female | RNAseq |
| 15505 | PCW12 | rsa(13,15,16,18,21,22,X)x2 | Female | |
| 15478 | PCW12 | rsa(13,15,16,18,21,22)x2,(X,Y)x1 | Male | |
| 15165 | PCW16 | rsa(13,15,16,18,21,22,X)x2 | Female | |
| 15413 | PCW16 | rsa(13,15,16,18,21,22)x2,(X,Y)x1 | Male | |
| 15513 | PCW16 | rsa(13,15,16,18,21,22,X)x2 | Female | |
| 15014 | PCW20 | rsa(13,15,16,18,21,22,X)x2 | Female | |
| 15060 | PCW20 | rsa(13,15,16,18,21,22)x2,(X,Y)x1 | Male | |
| 15102 | PCW20 | rsa(13,15,16,18,21,22,X)x2 | Female | |
| 14794 | PCW12 | rsa(13,15,16,18,21,22,X)x2 | Female | Live imaging |
| 15546 | PCW12 | rsa(13,15,16,18,21,22,X)x2 | Female | |
| 15316 | PCW12 | rsa(13,15,16,18,21,22,X)x2 | Female | |
| 15478 | PCW12 | rsa(13,15,16,18,21,22)x2,(X,Y)x1 | Male | |
| 15483 | PCW16 | rsa(13,15,16,18,21,22)x2,(X,Y)x1 | Male | |
| 15165 | PCW16 | rsa(13,15,16,18,21,22,X)x2 | Female | |
| 15314 | PCW16 | rsa(13,15,16,18,21,22)x2,(X,Y)x1 | Male | |
| 15413 | PCW16 | rsa(13,15,16,18,21,22)x2,(X,Y)x1 | Male | |
| 14787 | PCW20 | rsa(13,15,16,18,21,22)x2,(X,Y)x1 | Male | |
| 15383 | PCW20 | rsa(13,15,16,18,21,22)x2,(X,Y)x1 | Male | |
| 15048 | PCW20 | rsa(13,15,16,18,21,22)x2,(X,Y)x1 | Male | |
| 15060 | PCW20 | rsa(13,15,16,18,21,22)x2,(X,Y)x1 | Male | |
| 14807 | PCW12 | rsa(13,15,16,18,21,22,X)x2 | Female | Immunohistochemical quantification |
| 14794 | PCW12 | rsa(13,15,16,18,21,22,X)x2 | Female | |
| 15316 | PCW12 | rsa(13,15,16,18,21,22,X)x2 | Female | |
| 15413 | PCW16 | rsa(13,15,16,18,21,22)x2,(X,Y)x1 | Male | |
| 15314 | PCW16 | rsa(13,15,16,18,21,22)x2,(X,Y)x1 | Male | |
| 15130 | PCW16 | rsa(13,15,16,18,21,22,X)x2 | Female | |
| 15027 | PCW20 | rsa(13,15,16,18,21,22,X)x2 | Female | |
| 15102 | PCW20 | rsa(13,15,16,18,21,22,X)x2 | Female | |
| 15048 | PCW20 | rsa(13,15,16,18,21,22)x2,(X,Y)x1 | Male | |

**Supplementary methods table 2: primary and secondary antibodies**

| Primary antibodies | | | | |
|---|---|---|---|---|
| **Protein Target** | **Host Species** | **Concentration** | **Supplier** | **Emission Wavelength** |
| TUBB3 | Rabbit | 1:500 | Covalence | N/A |
| TUBB3 | Mouse | 1:500 | Covalence | N/A |
| SM22 | Rabbit | 1:500 | Abcam | N/A |
| ITPR1 | Rabbit | 1:500 | Alomone Labs | N/A |
| ADORA2B | Rabbit | 1:250 | Alomone Labs | N/A |
| P2RX2 | Rabbit | 1:500 | Alomone Labs | N/A |
| CACNA1C | Mouse | 1:250 | Abcam | N/A |
| SCFR | Goat | 1:500 | R&D Systems | N/A |
| ANO1 (TMEM16a) | Rabbit | 1:100 | | N/A |
| **Secondary antibodies** | | | | |

| Protein Target | Host Species | Concentration | Supplier | Emission Wavelength |
|---|---|---|---|---|
| Rabbit | Goat | 1:500 | Invitrogen | 488 |
| Rabbit | Goat | 1:500 | Invitrogen | 568 |
| Mouse | Goat | 1:500 | Invitrogen | 488 |
| Mouse | Goat | 1:500 | Invitrogen | 568 |
| Nuclei (DAPI) | N/A | 1:1000 | Sigma Aldrich | 350 |

**Supplementary data table 1: Expression changes in calcium and purinergic pathway-related genes in the small intestine between PCW12 and PCW20**

| Calcium pathway-related genes | | |
|---|---|---|
| **Gene** | **logFC: PCW16 vs PCW12** | **logFC: PCW20 vs PCW12** |
| ITPR1 | 0.60 | 0.24 |
| ITPR2 | 0.41 | 0.46 |
| PLCB2 | 0.77 | 1.02 |
| PLCB4 | 0.75 | 1.02 |
| CACNA1C | 0.49 | 0.37 |
| RYR1 | 0.64 | 0.58 |
| RYR3 | 0.63 | 0.25 |
| PRKCB | 0.92 | 1.24 |
| P2RX1 | 2.13 | 2.29 |
| P2RX7 | 0.68 | 0.75 |
| ADORA2A | 0.40 | -1.69 |
| **Purinergic pathway-related genes** | | |
| **Gene** | **logFC: PCW16 vs PCW12** | **logFC: PCW20 vs PCW12** |
| P2RX1 | 2.13 | 2.29 |
| P2RX2 | 0.66 | 1.49 |
| P2RX7 | 0.68 | 0.75 |
| P2RY13 | 0.74 | 1.53 |
| P2RY10 | 2.35 | 2.29 |
| P2RY4 | 1.13 | 1.48 |
| ADORA2B | 0.32 | 1.15 |
| ADORA3 | 0.43 | 1.49 |

**Contractility Analysis: ST Map construction.**

Movies of small intestine motility were imported into ImageJ (FIJI), spatially cropped and saved as multi-slice tiff stacks. The spatial pixel dimension was calculated from a small ruler placed in the field of view (FOV, **Supp. Fig. 4A**). Multi-slice tiff stacks were imported into custom software (Volumetry G9j: GWH) where the following routines were applied:

a) Image sequences were calibrated for time and space
b) Bit depth changed from 8-bit to 16-bit to reduce posterization after filtering
c) Frame averaging (±1 frame) and spatial smoothing (Gaussian 3x3x0.6SD or 5x5x1SD) were used to reduce granular noise
d) An intensity threshold was used to demarcate intestinal segments from the black background followed by conversion of intestinal segments to a coordinate-based particle format before final assembly into a 4D spatio-temporal object (4DSTO: x=length, y=diameter, z=time and colour intensity=grayscale: **Supp. Fig. 4B**: see Longden et al., 2021[1])
e) The outer perimeter of the intestine was located (2D Marching Cubes algorithm) and the midline spline calculated and smoothed for each slice of the 4DSTO (±5 pixel average: **Supp. Fig. 4C, Supplementary Movie #1**)
f) The tangent of the midline spline was calculated at every pixel along the preparation and the distance from the midline spline to the top and bottom perimeters, perpendicular to the spline, was measured to calculate the intestinal diameter (**Supp. Fig. 4C, inset**).
g) The diameters of the preparation at every pixel along the segment were converted to a color-coded horizontal line (grayscale-graded according to diameters), with every line from subsequent frames of the movie (extracted from the 4DSTO) stacked from top to bottom as a spatiotemporal map (ST Map (diameter: ); **Supp. Fig. 4D**). Time is represented on the y-axis (top=start or recording) and distance along the preparation represented on the x-axis (left=oral end of the preparation), with the level of grayscale denoting diameter (whiter=more dilated).

**Contractility Analysis: ST Map analysis**

ST maps of intestinal segments showed rhythmic contractility visualised as alternating dark (reduced intensity/diameter: contractions) and light (increased intensity/diameter: bulges) bands that propagated in the oral (sloping to the left) or anal (sloping to the right) direction (**Supp. Fig. 4D**).

To isolate only dynamic contractions, variations in background diameter due to different amounts of contents in the preparation were filtered using a long span moving maxima (20s timespan). The resulting maxima map (**Supp. Fig. 4E**) was subtracted from the original ST map to: i) isolate only dynamic contractions and ii) assess volume relocation changes during recordings.

Dynamic contractions were further enhanced by using a recursive peak detection algorithm that used a moving 3-point (triad) sub-sample arrayed over a series of timespans to detect inflection points corresponding to diameter minima (maximum contraction). The accumulation of marked peaks identified in the series of timespans (5.5-16.5 seconds, **Supp. Fig. 4F**), was thresholded (>50% cumulative intensity, **Supp. Fig. 4G**) and skeletonized (**Supp. Fig. 4H**) to demarcate the spatio-temporal trajectory of contractions. Results were saved as spatio temporal (ST) contraction filaments.

The contraction trajectories were used to calculate frequencies, intervals, velocities, diameters and the pattern of initiation sites and collision sites (**Supp. Fig. 4I-M**). Following filtering to remove spurious noise (trajectories < 10 pixels [~0.1mm] in length), the time interval between paired contraction trajectories was calculated for all points in which they occupied the same position along the preparation. The proportion of the recording that was contracting at a particular frequency (interval) was calculated and presented as a histogram. Intervals were color-coded to construct frequency (interval) maps (FMaps: see Hennig et al., 2010[2]) to visualize dynamic changes in frequency during recordings (**Supp. Fig. 4J**). In a similar manner, instantaneous velocities were calculated from the slope of contraction trajectories at every point (spatial span ± 5 pixels [~0.05mm]) to construct velocity maps (VMaps: **Supp. Fig. 4K**).

Propagating contractions travel in both directions away from an initiation site. Multiple initiation sites along a preparation result in collisions between contractions travelling toward each other ("V-shaped" ST contraction filaments in **Supp. Fig. 4I** and blue->red transitions in **4K**), preventing propagation of contractions over long distances. The number of regions displaying coherent propagations, called "directional domains", along each preparation was quantified by summing the directions of propagation for every point along the preparation from ST filaments used to construct velocity maps (spatial span ± 0.75mm: **Supp. Fig. 4L**). We chose to measure the distance of uni-directional domains instead of bi-directional domains as initiation sites were often located on the extreme ends of the preparation where contractions could only propagate in one direction. The number of uni-directional domains (in between initiation and collision sites) was counted, with transitional areas receiving a half score (0.5). Results were normalized to a standard preparation length of 30 mm.

**RNAseq analysis**

Raw reads initially underwent QC and trimming with the program fastp. Then, high-quality, trimmed reads were mapped on the human genome (version hg38) with the mapper STAR, using default options. Raw counts were then generated using the program featureCounts. The count table was imported in R/v4.2.1[3], filtered to include genes with more than 5 reads in at least 70% of the samples (removing uninformative genes, i.e., genes with low expression level that would contribute to background noise[4]) and normalised using the RUVseq/v1.30.0[5] library with "k=2" factors of unwanted variation. Differential gene expression was assessed with the library edgeR/v3.38.4[6] using a Generalized Linear Model, following the recommendations on the edgeR user's guide, in a one-way ANOVA test where samples of PCW12 were used as baseline and genes that differ between any of the developmental conditions (i.e., PCW12, PCW16 and PCW20) were identified. Genes having an adjusted p-value (i.e., fold discovery rate, FDR) lower than 0.05 were deemed significant.

Functional analysis was conducted using the clusterProfiler/v4.4.4[7] package by performing Over-Representation Analysis (ORA) with the function “enricher” and Gene Set Enrichment Analysis[8] (GSEA) with the function “gsea”. Functional databases were retrieved from the gProfiler’s website (https://biit.cs.ut.ee/gprofiler/gost) and pathways with adjusted p-value (i.e., FDR) of ≤ 0.2 were deemed significant.

**Quantification of immunofluorescent staining**

Images for quantification were captured on an LSM710 confocal microscope (Zeiss, Germany) using a 40X (NA1.2) water immersion objective (Zeiss, Germany). For each protein target/age quantification was conducted on n=3 samples, with 2 sections examined per sample. 4 representative images (512 X 512 pixels) of the intestinal musculature were collected from each section (for a total of 8 images per patient sample, and 36 images per age under examination) in a cross-shaped array at 63X magnification. Images were imported into FIJI software[9] and quantified in a blinded fashion. Briefly, images were converted into 8-bit format and the *Despeckle* function applied followed by *Threshold* (default setting used). Each protein of interest was normalised against their respective DAPI images and expressed as a percentage. The results from the 8 images of 2 sections per patient sample were averaged.
